# A relative comparison between Hidden Markov- and Log-Linear-based models for differential expression analysis in a real time course RNA sequencing data

**DOI:** 10.1101/448886

**Authors:** Fatemeh Gholizadeh, Zahra Salehi, Ali Mohammad banaei-Moghaddam, Abbas Rahimi Foroushani, Kaveh kavousi

## Abstract

With the advent of the Next Generation Sequencing technologies, RNA-seq has become known as an optimal approach for studying gene expression profiling. Particularly, time course RNA-seq differential expression analysis has been used in many studies to identify candidate genes. However, applying a statistical method to efficiently identify differentially expressed genes (DEGs) in time course studies is challenging due to inherent characteristics of such data including correlation and dependencies over time. Here we aim to relatively compare EBSeq-HMM, a Hidden Markov-based model, with multiDE, a Log-Linear-based model, in a real time course RNA sequencing data. In order to conduct the comparison, common DEGs detected by edgeR, DESeq2 and Voom (referred to as Benchmark DEGs) were utilized as a measure. Each of the two models were compared using different normalization methods. The findings revealed that multiDE identified more Benchmark DEGs and showed a higher agreement with them than EBSeq-HMM. Furthermore, multiDE and EBSeq-HMM displayed their best performance using TMM and Upper-Quartile normalization methods, respectively.

## Introduction

The information stored in a single gene is transcribed into RNA and then translated into protein(s). The set of RNAs transcribed at a given time or developmental stage, known as transcriptome, reveals the state of a cell and can be used to study gene expression (1). Next Generation Sequencing (NGS) technologies have made it plausible to sequence RNA transcripts in a high-throughput manner. The method developed for this purpose is known as RNA sequencing (RNA-seq) (2).

Whatever NGS platform for RNA-seq is chosen, which depends on the experimental goals, millions of sequence reads will be provided as output (3). The number of reads mapped to a gene or transcript (read counts) can be obtained as the estimate of the gene or transcript’s expression level (4, 5). There are a number of tools available to obtain read counts, such as featureCounts (6) and HTSeq-count (7).

Identification of differentially expressed genes (DEGs) has attracted attention in RNA-seq studies due to its application in relating genotype and phenotype to each other and detecting pathological mechanisms underlying diseases (8).

The statistical methods for detection of DEGs are either non-parametric or parametric. Many parametric methods assume that gene expression values follow a Negative Binomial (NB) distribution (9). Most of available tools for differential expression analysis detect a gene as DE if its expression changes in at least one condition. However, in time course experiments where researchers are interested in genes that change over time such as disease stages, these methods do not account for correlation and dependencies between two consecutive time points (10). Therefore, their analysis power is likely to decrease (11). Until now, many attempts have been made to surmount these challenges in time course studies and a number of tools have been developed and implemented for this purpose (10, 12–20). Yet, no standard method has been established for time course differential analysis (11).

In this study, our analysis is focused on a relative comparison of two approaches in detection of DEGs in a real time course study, as the performance of these approaches has not been compared to each other before. The first approach implemented by EBSeq-HMM tool (10) is based on a Hidden Markov Model (HMM) and is particularly designed for ordered RNA-seq data such as in time course experiments. The second approach is implemented by multiDE (21) and works based on a log-linear model. This tool has been designed for experiments with multiple treatment conditions but is capable of accounting for correlation between matched samples in different conditions. To compare these tools, the common DEGs found by three benchmark methods: edgeR(22), DESeq2(23) and Voom(24) were used as a gold standard measure. The two models were compared by different normalization methods. Moreover, models’ false positives were investigated using a reference set of constant housekeeping genes.

## Materials and methods

### Statistical Models

In the following, we briefly describe the main points of the two statistical models, EBSeqHMM and multiDE. More elaborated description of the models is accessible in the original papers (10, 21).

**EBSeq-HMM** is an R bioconductor package (available at: https://bioconductor.org) allowing detection of DEGs in time course RNA-seq experiments. It assumes a NB distribution for gene expression and requires at least three time points. Genes are divided into three categories: dynamic, sporadic and constant genes.

If *X*_*gti*_ denotes gene expression estimation for gene *g* at time *t* in sample *i* (*i* = 1,…, *n*_*t*_, where *n*_*t*_ is the number of samples at time t), and *μ*_*gt*_ denotes the latent mean expression for gene *g* at time *t*, three states can describe changes in *μ*_*gt*_ between each two time points *t-1* and *t*: state 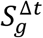 as *UP* if *μ*_*g*,*t*_, state 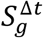 as *DOWN* if *μ*_*g*–*t*_ > *μ*_*g*,*t*_ and state 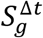 as *EE* (Equally Expressed) if *μ*_*g*–*t*_ > *μ*_*g*,*t*_. Then, 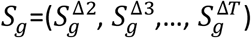 can be obtained to estimate the most likely expression path for a gene over time.

In time course studies, expression value for gene *g* at time *t*, *X*_*gt*_, not only depends on the one at previous time, *X*_*g*,*t*–1_, but also on state 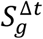. Dynamic genes experience continuous changes in their expression values as time passes and therefore each state 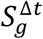 depends on the former state 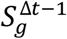. Hence, the state process is assumed to be described by a Markov chain. The emission model at each time is assumed to be Beta-NB as obtained in the original paper (10). For constant and sporadic genes, which do not represent continuous changes over time, states are assumed to be independent from each other. In this case, the emission model remains the same but the state process is not assumed to be explained by a Markov chain. To accommodate dependence among gene expressions, an auto-regressive approach is used. It means that, for dynamic genes, an auto-regressive hidden Markov approach is used while sporadic and constant genes are handled using an auto-regressive non-hidden Markov approach. Since the type of expression path is not known beforehand, a two-component mixture model is held over all types of genes.

To detect DEGs, EBSeq-HMM estimates the posterior probability (PP) of each possible expression path for a gene (*PP*(*S*_*g*_ = *s*|*X*_*g*_)). Under a predefined FDR (False Discovery Rate) equal to *α*, a gene is said to be DE if its PP of being constant is less than or equal to *α*. To classify differentially expressed genes into their most likely expression path, the path with PP≥0.5 is chosen. EBSeq-HMM has two normalization methods available in its package, denoted by Median-of-Ratios (default) and Upper-Quartile.

**MultiDE** is an R package (available at: http://homepage.fudan.edu.cn/zhangh/softwares/multiDE) developed for differential expression analysis in experiments with multiple treatment conditions and needs a minimum of two biological replicates in each condition. This model can be used for either matched samples (to decrease bias due to confounding) or unmatched samples. To identify DEGs, a log linear model consisting of two main effects (gene and condition) and an interaction effect (gene*condition) which measures the relative association between gene and condition is used. Since the current study is time course, we assume the situation is matched and treatment conditions are the time points. Read counts are assumed to follow a NB distribution.

Assume *X*_*gti*_ as described in EBSeq-HMM model. If *X*_*gti*_ follows a NB distribution with expectation *μ*_*gt*_ and dispersion parameter *φ*_*g*_, then the model is defined as:

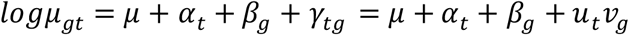

Where *μ* is the grand mean and *α*_*t*_ and *β*_*g*_ are the main effects for time *t* and gene *g*. The interaction effect between time *t* and gene *g* is denoted by *γ*_*tg*_:= *u*_*t*_*v*_*g*_ which measures the relative association between time *t* and expression value of gene *g*. After some restrictions and calculation provided in detail in the original paper, it can be resulted that if at least one DEG exists between *t* time points, then for at least one gene, *γ*_*tg*_ ≠ 0 is held. Therefor, *u*_*t*_ ≠ 0 for at least one *t* under *γ*_*tg*_:= *u*_*t*_*v*_*g*_. When both *γ*_*tg*_ ≠ 0 and *u*_*t*_ ≠ 0, then *v*_*g*_ can measure the relative association. Therefore, the null hypothesis to detect whether gene *g* is differentially expressed between *t* time points is defined as *H*_*g*_: *v*_*g*_ = 0. The Wald test statistic is 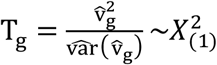 under null hypothesis. To learn how to estimate the parameters, we refer the readers to the original paper (21).

It must be noted that four normalization methods implemented in multiDE package are denoted by MEDIAN (default), TOTAL, QUANTILE and TMM.

### Real Data

The dataset comes from a time course study by Piras et al. (2017) to determine the consequence of lentiviral transduction of human HSPC. They examined the transcriptional profiling of bone marrow derived CD34+ cells transduced with different lentiviral vectors at different time points. (25). The table of raw read counts was downloaded from NCBI GEO database (GEO accession ID: GSE92652). The sub table of read counts for cells transduced with purified LV was used for the analysis. Data were collected in 5 time points with three replicates each.

### Significance Level

Significance level for all differential expression analyses was 0.05.

## Results

To filter out lowly expressed genes, genes without a minimum of 10 read counts in at least 70 percent of samples were ignored and discarded. Afterwards, there remained 17746 genes out of 55386 genes. EBSeq-HMM and multiDE were compared by different normalization methods they had implemented in their packages but the normalization method in the benchmark models remained by default. To identify differentially expressed genes, all 5 models were run by their default settings except for EBSeq-HMM in *EBSeqHMMTest* command where we set Qtrm and QtrmCut to 100 and 0, respectively. Whenever needed, P-values were adjusted using Benjamini–Hochberg (BH) method (26).

### Number of DEGs detected

Using Median-of-Ratios normalization, EBSeq-HMM lead to detection of 15250 DEGs (85.93% of total genes) while by the use of Upper-Quartile normalization, EBSeq-HMM detected 15224 DEGs (85.79% of total genes). Although the findings seem comparable, the number of detected DEGs using either of the two normalization methods was too large so that we only took confident DEGs (DEGs with PP of most likely expression path ≥ 0.5) for further analysis.

EBSeq-HMM (Median-of-Ratios), EBSeq-HMM (Upper-Quartile), multiDE (MEDIAN), multiDE (QUANTILE), multiDE (TMM), multiDE (TOTAL), identified 4336, 4336, 7457, 7326, 7995, 7296 DEGs, respectively (Fig. 1).

**Fig.1.**
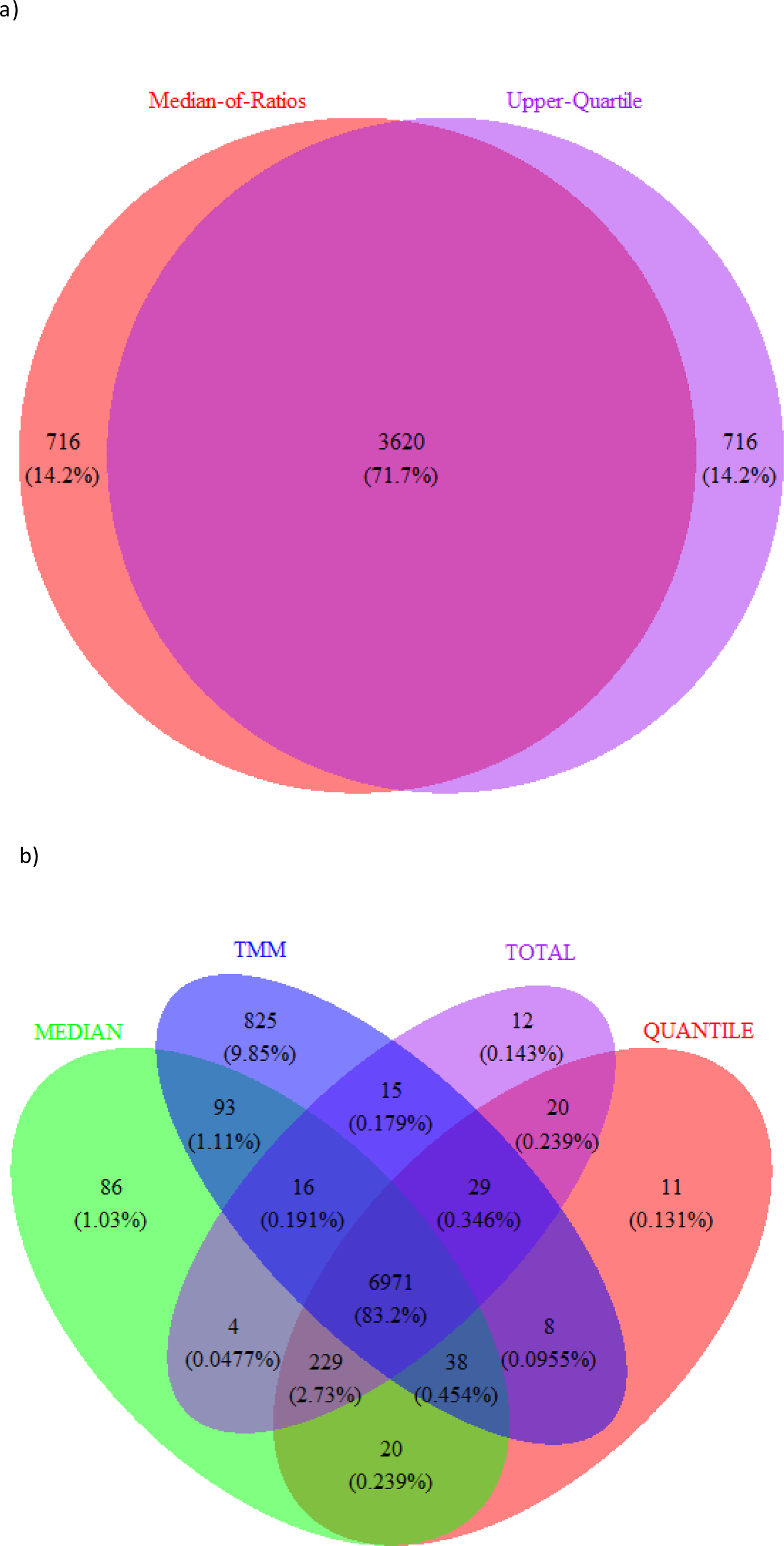
Number of DEGs detected by different normalization methods. a) EBSeq-HMM model; b) multiDE model.

### Models Agreement with Benchmark DEGs

EdgeR, DESeq2 and Voom, each found 7858, 8065, 5929 DEGs, respectively. The number of DEGs overlapped by all three models was 5828. We refer to these genes as “Benchmark DEGs” and used them as a measure to compare EBSeq-HMM with multiDE in terms of the proportion of Benchmark DEGs they could find and the percentage of their agreement with Benchmark DEGs. Table 1 reveals that despite the fact that Median-of-Ratios and Upper-Quartile normalization methods resulted in the same number of DEGs using EBSeq-HMM, Upper-Quartile normalization lead to identification of a slightly more Benchmark DEGs (34.09%). On the other hand, by the use of TMM normalization method, multiDE could identify the largest proportion of Benchmark DEGs (74.74%). Using any of the normalization methods, multiDE was able to find more than twice Benchmark DEGs as EBSeq-HMM. Considering the default normalization methods, the common and unique numbers of Benchmark DEGs detected by EBSeq-HMM and multiDE are shown in Fig. 2. It is easy to see that multiDE detected more novel DEGs (2750 genes). On the other hand, this tool displayed more agreement with Benchmark DEGs than EBSeq-HMM did. 57.5% of the genes detected as DE by multiDE were Benchmark while 45.64% genes declared as DE by EBSeq-HMM were Benchmark (table 1).

**Table 1.** A comparison of EBSeq-HMM and multiDE using different normalization methods

| **Model** | **Number of DEGs detected** | **Model's agreement with Benchmark DEGs (%)** | **Benchmark DEGs found by model (%)** |
| --- | --- | --- | --- |
| EBSeq-HMM (Median-of-Ratios) | 4336 | 45.64 | 33.96 |
| EBSeq-HMM (Upper-Quartile) | 4336 | 45.83 | 34.09 |
| multiDE (MEDIAN) | 7457 | 57.50 | 73.58 |
| multiDE (QUANTILE) | 7326 | 58.07 | 72.99 |
| multiDE (TMM) | 7995 | 54.48 | 74.74 |
| multiDE (TOTAL) | 7296 | 58.14 | 72.79 |

**Fig. 2.**
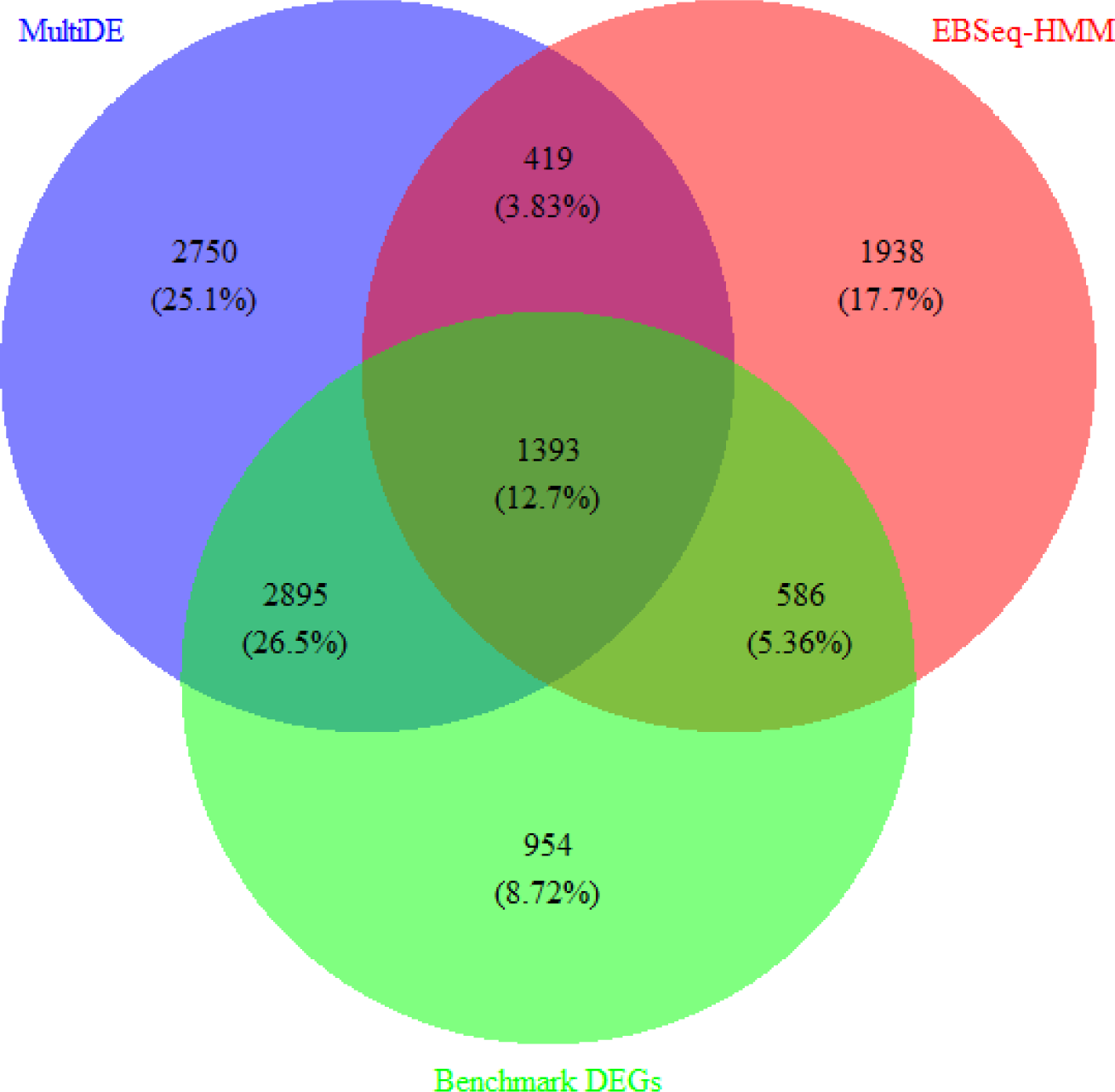
Number (percentage) of novel and common DEGs found by EBSeq-HMM and multiDE.

To further evaluate EBSeq-HMM, we divided DEGs with PP of most likely expression path more than 0.7 into three categories as shown in Table 2. It can be seen that 71.21% of the most confident genes (PP≥0.9) were Benchmark which accounted for 4% of all Benchmark DEGs. As the PPs decreased, model’s agreement with Benchmark DEGs decreased.

**Table 2.**
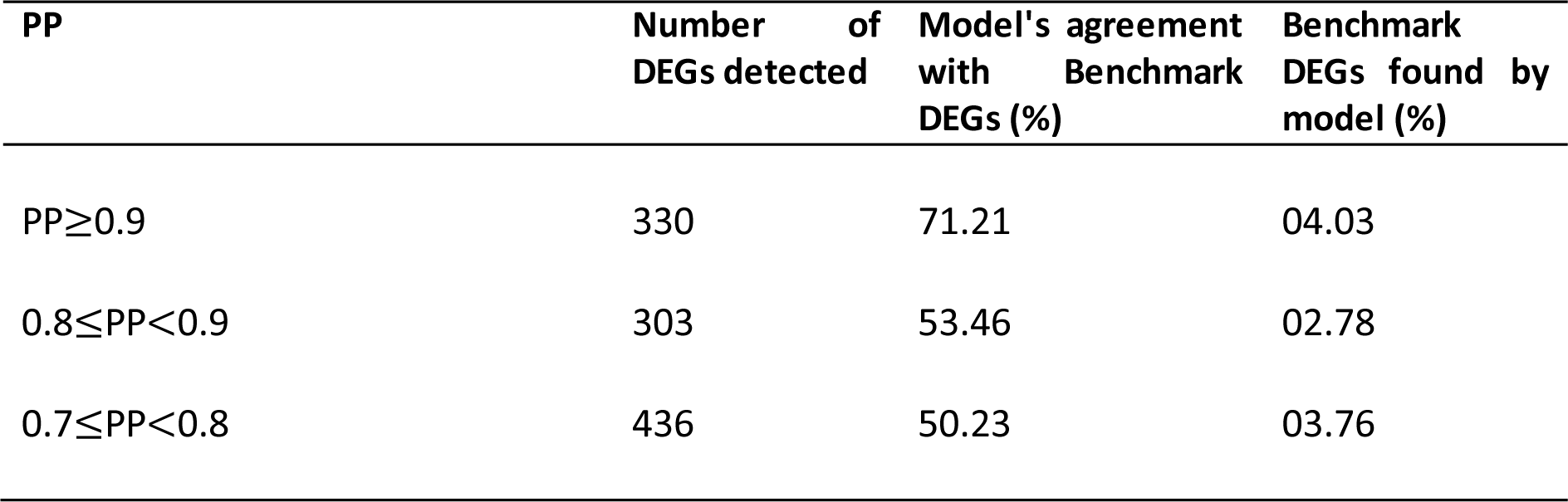
A comparison of DEGs identified by EBSeq-HMM based on PPs

We, then, made three categories of the adjusted p-values for DEGs found by multiDE (Table 3). Among them, the category of the most significant genes (P-value≤0.0001) had the highest agreement with Benchmark DEGs and a total of 42.86% of Benchmark DEGs were common with them. Whereas, the least significant genes (0.006<P-value<0.05) had the least agreement with Benchmark DEGs. They were able to detect only 10.84% of Benchmark DEGs.

**Table 3.** A comparison of DEGs found by multiDE based on P-values

| P-value | Number of DEGs detected | Model's agreement with Benchmark DEGs (%) | Benchmark DEGs found by model (%) |
| --- | --- | --- | --- |
| $P\text{-value} \leq 0.0001$ | 2870 | 87.04 | 42.86 |
| $0.0001 < P\text{-value} \leq 0.006$ | 2177 | 53.19 | 19.87 |
| $0.006 < P\text{-value} < 0.05$ | 2410 | 26.22 | 10.84 |

### Models’ False Positives

A housekeeping gene that is known to have a constant/stable expression in a cell across different treatments can be used as a reference to evaluate the models’ false positives. Eight genes namely *RPL41*, *RPLP0*, *RPS27*, *TUBA1B*, *RPSA*, *SLC25A3*, *ACTG1* and *EEF1G* have been introduced as a possible reference in gene expression studies of human bone marrow (27). *RPLP0*,*RPS27* were on the list of Benchmark DEGs and *EEF1G* did not have a minimum of 10 read counts in at least 70 percent of samples, so that we excluded them from the study. A glance at Table 4 reveals that EBSeq-HMM detected none of the housekeeping genes as DE while multiDE had 2 to 3 false positives depending on the normalization methods.

**Table 4.** Evaluation of false positives of models using housekeeping reference genes

| model | *RPL41* | *TUBA1B* | *RPSA* | *SLC25A3* | *ACTG1* |
| --- | --- | --- | --- | --- | --- |
| EBSeq-HMM (Median-of-Ratios) | × | × | × | × | × |
| EBSeq-HMM (Upper-Quartile) | × | × | × | × | × |
| multiDE (MEDIAN) | ✓<br>P-value=0.015 | × | ✓<br>P-value=0.014 | ✓<br>P-value=0.042 | × |
| multiDE (QUANTILE) | ✓<br>P-value=0.021 | × | ✓<br>P-value=0.019 | × | × |
| multiDE (TMM) | ✓<br>P-value=0.005 | × | ✓<br>P-value=0.005 | ✓<br>P-value=0.029 | × |
| multiDE (TOTAL) | ✓<br>P-value=0.021 | × | ✓<br>P-value=0.017 | × | × |

X: The gene was not detected as DE. ✓: The gene was detected as DE. P-values are given for DEGs.

## Discussion

By now, a number of studies have made attempts to compare statistical methods for differential expression analysis in RNA-seq time course experiments. However they have generally focused on methods developed specifically for time course data (11, 28). Those studies did not take it into account that other methods developed for multiple treatment conditions and pairwise comparisons might be surprisingly able to detect DEGs in a more efficient way. Therefore, such methods did not receive enough attention in their studies.

In this study, we evaluated the relative performance of two Hidden Markov- and Log-Linear-based statistical models in detection of DE genes in a real time course RNA-seq data. The Hidden Markov-based model, EBSeq-HMM, was particularly developed for time course experiments while the log-linear based model, multiDE, was proposed for multiple treatment conditions.

Findings revealed that EBSeq-HMM detected a high percentage of genes (more than 85% of all genes) as DE which indicated its poor performance. As said by a previous study (11), that is because the constant component of the model assumes a gene’s expression to be exactly equal between two time points while this is improbable to happen in real situations. In order to have a reasonable number of DEGs to conduct the comparison, we had to put a limit on the results and focus our study on the confident DEGs. Compared to multiDE, EBSeq-HMM detected less Benchmark DEGs and had a lower agreement with them even though its confident DEGs were selected for analysis. Furthermore, multiDE’s better performance was resulted while this model was in fact designed for multiple treatment conditions not time course experiments.

It is interesting to note that different normalization methods in each model performed comparably. Yet, Upper-Quartile normalization in EBSeq-HMM and TMM normalization in multiDE lead to detection of slightly more Benchmark DEGs. The latter is consistent with the findings of the authors of multiDE (21). They suggested that TMM normalization method performed better than the other three normalizations available in multiDE package and suggested using TMM in real data.

In terms of false positives, multiDE underperformed EBSeq-HMM. This can be caused by the fact that we limited the results of EBSeq-HMM as already mentioned. Taking into account all DEGs detected by EBSeq-HMM could potentially lead to more false positives.

Finally, one drawback to multiDE was that its results may not be used directly as further analysis was required to adjust p-values for multiple comparison which was not implemented in multiDE package.

Overall, multiDE detected more Benchmark DEGs and had a higher level of agreement with them than EBSeq-HMM. However, EBSeq-HMM is capable of assigning each gene to their most likely expression path which can be beneficial when one needs to know the expression path of a given gene over time. To obtain better results, it is recommended to run multiDE and EBSeq-HMM using TMM and Upper-Quartile normalization methods, respectively, as they displayed their best performance using these normalization methods. This study showed that, even though multiDE was developed for multiple treatment conditions and was not able to accommodate dependencies overtime, it outperformed EBSeq-HMM, which was specifically designed for time course experiments. It should be taken into account that all of the above results have been obtained on a real data and therefore, they cannot be generalized to other situations and data confidently.

## Acknowledgement

We would like to thank School of Public Health at Tehran University of Medical Sciences, Institute of Biochemistry and Biophysics (IBB) at University of Tehran and Clinical Research Development Center of Persian Gulf Martyrs Hospital for supporting us and the comments they made to improve this paper.

